# Extracting generator-specific information from irregular brain field potentials: a comparative study of feature-based and deep learning approaches

**DOI:** 10.64898/2026.11.27.769679

**Authors:** Nicusor Cosmin-Toader, Ana Ramírez-Frías, Ricardo Muñoz-Arnaiz, Juan Bautista-Aballay, Faezeh Ashouri, Julia Makarova, Marcos García-Lorenzo, Oscar Herreras

**Author notes:** Correspondence: Nicusor Cosmin-Toader, VG-LAB, Universidad Rey Juan Carlos, Móstoles, Spain, or, Oscar Herreras, Centro de Neurociencias Cajal, CNC-CSIC, Av. León 1, Alcalá de Henares, 28805, Spain. Co-first authors (Nicusor Cosmin-Toader, Ana Ramírez-Frías, Ricardo Muñoz-Arnaiz). Co-senior authors (Marcos García-Lorenzo, Oscar Herreras).

## Abstract

Brain field potentials (FPs) reflect the combined activity of multiple neural sources, challenging the accuracy of their temporal series and their spatial origin difficult to establish from waveform inspection alone. However, FP generators obtained by independent component analysis exhibit characteristic temporal patterns and unique spatial profiles that allow their identification, even during apparently irregular activity. Here, we compared feature-based machine learning (ML) and end-to-end deep learning (DL) approaches to assess how effectively this information can be extracted from short segments and whether learning directly from raw signals offers advantages over predefined signal descriptors. We analyzed irregular FP-generator activity from multiple hippocampal generators in a dataset pooled across experimenters, recording devices, and electrode types. Feature-based approaches used predefined spectral and nonlinear descriptors to construct multidimensional representations, which were classified using seven ML algorithms: bagged decision trees (BAG), light gradient boosting machine (LGBM), quadratic discriminant analysis (QDA), random forest (RF), support vector machines (SVM), extreme gradient boosting (XGB) and multilayer perceptron (MLP). In parallel, convolutional neural networks (CNNs) and transformer architectures learned discriminative representations directly from raw signals. Performance was evaluated using animal-independent data partitions to assess generalization to previously unseen animals. Both approaches discriminated among FP generators from short segments of irregular activity. Raw-signal models generally achieved higher classification performance than feature-based approaches, although the latter also captured substantial generator-specific information. This suggests that discriminative information extends beyond the predefined descriptors and includes temporal patterns that can be exploited through direct signal learning. Classification across animals further supports the reproducibility of generator-related information despite variability in experimental conditions and recording procedures. These findings provide a scalable quantitative approach to identifying FP generators from irregular activity, opening new possibilities for investigating electrophysiological signals across physiological and pathological conditions.

---

**Keywords**: Field potentials, Irregular activity, feature-based descriptors, machine learning, deep learning, nonlinear signal analysis

## 1. Introduction

Mathematical and computational models provide powerful tools for extracting information from complex signals, including brain EEG and field potentials (FPs). Traditional approaches have focused primarily on recognizable patterns as markers of behavioral tasks, cognitive states, and pathological conditions (Nunez and Srinivasan 2006; Buzsáki et al., 2012). However, these patterns account for only a small fraction of recorded FP activity, which predominantly consists of irregular fluctuations whose physiological significance remains difficult to establish (Eckhorn, 1994; Bullock et al., 2003; Monto et al., 2008). This reflects both the limitations of analytical approaches for characterizing such activity and the difficulty of relating measured signal properties to their underlying neural sources (Herreras, 2016), leaving much of the potentially informative signal unexplored.

Interpreting irregular FP activity is challenging because recorded signals reflect the superposition of contributions from large numbers of microscopic neural sources, mostly synaptic currents, whose recruitment and relative contributions vary over time and space (Lorente de Nó, 1947). Different source configurations can produce similar extracellular potential patterns, making the inverse problem of inferring the underlying sources inaccessible (Pascual-Marqui, 1996). For intracranial recordings, multisite sampling combined with spatial decomposition techniques, such as independent component analysis (ICA), has substantially alleviated this problem (Makarov et al., 2010). ICA separates multisite raw FPs into a limited number of spatially coherent components, termed FP generators, identified in both humans and rodents (Benito et al., 2014; Schomburg et al., 2014; Orellana et al., 2023; Makarova et al., 2025). In some rodent hippocampal generators, experimental evidence and computational models have further enabled their attribution to specific anatomical pathways with stable, anatomically supported geometries (Makarova et al., 2011; Martín-Vázquez et al., 2016; Torres et al., 2019). These components have also proved useful for exploring dynamical relationships between oscillatory activities and their associations with behavior (López-Madrona et al., 2022; Fernández-Ruiz et al., 2021, 2023). Importantly, FP generators provide an intermediate level of aggregation of microscopic contributions that is more tractable for quantitative analysis, avoiding the temporal degradation caused by the mutual distortion of mixing fields from distinct mesoscopic sources (Herreras et al., 2015; Herreras, 2016; Whitmore and Linn, 2016). Although their temporal activity remains predominantly irregular (Hernández-Recio et al., 2023), it can now be studied as an expression of the dynamics of spatially coherent neural populations rather than as undifferentiated background activity (López-Madrona et al., 2020, 2024; Nasretdinov et al., 2023).

Long time series of FPs or FP generators can be characterized using complementary measures, ranging from conventional statistical and spectral descriptors to fractal, entropy-based, and other nonlinear measures (Lau et al., 2022; Averna et al., 2023). In FP generators, combinations of these features provide a multivariate representation that may capture properties not readily apparent from visual inspection. Recognizable rhythms, including gamma, alpha, and theta oscillations, occur in several hippocampal FP generators (Benito et al., 2014, 2016; López-Madrona et al., 2020; Fernández-Ruiz et al., 2021, 2023; Hernández-Recio et al., 2023). However, much of their activity consists of irregular sequences of waves with varying amplitudes and durations, reflecting the stochastic and sparse nature of population dynamics (Pijn et al., 1991; Festa et al., 2021). Although the reliability and interpretation of some non-canonical measures remain debated, previous work in our lab suggests that some limitations affecting their application to raw FPs can be reduced when they are applied to spatially separated FP generators (Makarov et al., 2023).

An important open question is how much information in these dominant irregular fluctuations is captured by predefined features and whether additional discriminative information can be extracted directly from the signals. Feature-based models operate on metrics selected in advance, allowing classification to exploit established signal properties (Muñoz-Arnaiz et al., 2025). Their performance depends, however, on feature selection and on how models combine these metrics. Comparing models of different complexity can help establish whether generator discrimination relies on relatively simple relationships among features or requires more flexible nonlinear mappings. End-to-end deep-learning (DL) models, in contrast, learn representations directly from input signals and may capture discriminative patterns not explicitly encoded in handcrafted features (Craik et al., 2019; Cai et al., 2025; Goldstein et al., 2025). The relative advantages of these approaches, including the amount of data needed to achieve reliable performance, therefore remain empirical questions for these signals and tasks.

In the present study, we compare feature-based machine-learning (ML) models with raw-signal based DL models to investigate how much discriminative information can be extracted from the irregular temporal activity of multiple FP generators. We address four complementary questions: to what extent different generators can be discriminated from short signal segments; how much generator-specific information is captured by predefined features; whether increasing model complexity improves discrimination within this feature space; and whether end-to-end models exploit additional discriminative patterns directly from the signals. Finally, we assess generalization to previously unseen animals using models trained on data pooled across different experimenters, recording periods, and acquisition systems. Classification performance on held-out animals provides an empirical measure of how reliably generator-specific information can be extracted from irregular FP activity, while allowing for the possibility that some temporal segments contain little or no discriminative information.

## 2. Methods

### 2.1. Experimental datasets

The dataset comprised intracranial electrophysiological recordings obtained from 3-4 months old female Wistar rats in several previous studies (Benito et al., 2016; Montero-Atalaya et al., 2023; Makarova et al., 2026). The experiments were performed in accordance with EU directive (2010/63/UE), and Spanish (RD 53/2013) and local (Autonomous Community of Madrid, Order 4/8/1988) guidelines regarding the use of laboratory animals, and the experimental protocols were approved by the Research Committee of the Cajal Institute. Animal housing, preparatory and surgery details can be found in the respective studies.

The data pooled for the present study was obtained by different lab members over a period of 10 years. A total of 30 different animals was used. In eight individuals, recordings from both left and right hemispheres were used, albeit using different epochs. Basic experimental settings were homogeneous but potentially relevant differences are listed in Supplementary Table 1. It includes experimenter, animal age, implanted probes (brand, type, number of shanks, number of recording channels, electrode spacing), and acquisition hardware. Also, additional stimulating and recording electrodes or injecting pipettes, which did not participate in data gathering, could vary.

All recordings were performed under urethane anesthesia (1.2-1.5 g/Kg b.w.), with the animals fastened to a stereotaxic device. Multichannel silicon recording probes were aimed at the dorsal hippocampus and spanned the CA1 and the Dentate Gyrus (DG). Signals were acquired using MultiChannel Systems or Open Ephys acquisition systems, at an initial sampling frequency of 10-20 kHz. A silver chloride wire placed beneath the neck skin served as the reference.

Four subsets (A to D: Table 1) were made according to some of these experimental features and used to check for their potential impact on training models.

**Table 1.** Experimental subsets defined by combinations of variables potentially affecting FP recordings and subsequent analysis. A complete description of the experimental variables is provided in Supplementary Table 1.

| Subset | Age | Probes | Channels | Animals | Segments |
|---|---|---|---|---|---|
| A | 4-5 mo | 1 | 32 | 6 | 5100 |
| B | 3 mo | 1 | 32 | 9 | 26084 |
| C | 3 mo | 1 | 16 | 7 | 6464 |
| D | 3 mo | 2 | 16 | 8 | 23120 |

### 2.2. Signal processing and FP-generator extraction by Independent Component Analysis (ICA)

We used epochs recorded in control conditions, i.e. prior to any treatment or manipulation. The length varied between 4 and 300 min across series. During acquisition a high-pass digital filter was set at 0.5 Hz to remove slow transient artifacts, and the signals were resampled at 1-4 kHz to speed up analyses.

FP epochs showing sustained rhythmic patterns, such as hippocampal theta and highly regular cortical delta oscillations (slow-wave activity, SWA), were excluded. The latter are detectable in hippocampal recordings with substantial amplitude through volume conduction (Torres et al., 2019). Across all experimental series, a comparable level of anesthesia was maintained (Supplementary Table 1) that allowed the occurrence of a broad range of transient electrographic patterns intermediate between these two extremes. Continuous recordings were segmented by excluding episodes of rhythmic activity lasting longer than 5 s, while brief or intermittent episodes of theta and irregular SWA were retained. The remaining segments were concatenated to form analysis epochs, each of which was allowed to contain no more than three gaps. Stereotyped events associated with particular FP generators, such as CA1 sharp waves and dentate spikes, were not excluded (Bragin et al. 1995; Fernández-Ruiz et al. 2012; Santiago et al. 2024). Given their brief duration and sporadic occurrence, these events were unlikely to affect the overall performance of the ML/DL classification, which was based on large numbers of short segments sampled across epochs lasting many minutes.

The effectiveness of ICA is influenced by the statistical properties and structure of the input signals (Bell and Sejnowski 1995; Stone et al. 2002). For intracranial FPs recorded with linear multielectrode arrays, source separation is highly efficient as probes can be positioned within the spatial domain occupied by the underlying source generators, thus resolving distinct voltage gradients arising from different synaptic inputs, including those converging onto the same neuronal population (Makarov et al. 2010; Benito et al. 2014). Each ICA component provides a time course and a group of electrode coefficients that, when ordered, they reproduce a spatial profile across recording sites. Such profiles correspond to linear sampling of the tridimensional extension of the fields produced by each spatially stable mesoscopic source, then termed as FP-generators. In former studies they have been used along with anatomical data to infer the source location, polarity and other geometrical properties (Benito et al., 2014; Herreras et al. 2015; Martín-Vázquez et al., 2015). In the hippocampus, some have been shown to correspond to pathway-specific activities (Fernández-Ruiz et al., 2012; Benito et al., 2014).

For the present study original FP series were re-analyzed by only one person (JM) using the kernel density ICA algorithm (KDICA) (Chen 2006) implemented in MATLAB (available at http://blogs.mat.ucm.es/vmakarov/downloads). The resulting FP generators were inspected and identified according to their spatial profiles, allowing assignment to the corresponding anatomical regions (Benito et al., 2014). Four generators were selected for the present study for their stability across time and individuals: (a) Schaffer (CA3 input to the CA1 region); (b) lacunosum-moleculare (LM: as yet unidentified, possibly composed from 2-3 inputs in this stratum: Hernández-Recio et al., 2023); (c) Lateral Perforant Path (LPP: lateral entorhinal input to the DG); and (d) Remote (Rem, a volume-conducted generator arising mostly from cortical regions: Torres et al., 2019). These generators have been extensively characterized in previous studies (Benito et al., 2014, 2016; Torres et al., 2019).

FP generators were resampled to a common rate of 1 kHz using the resampling function implemented in the MNE Python library (Gramfort et al., 2013). This procedure uses frequency-domain filtering to prevent aliasing.

#### 2.2.1. Segmentation into signal windows

The FP generator time series were divided into non-overlapping windows of 5 s (see Figure 1). This window duration was selected as a commitment between temporal resolution and successful classification probability based on a preliminary analysis (Muñoz-Arnaiz et al., 2025). Windowing was performed ensuring that no segment contained samples separated by an excluded interval. When an epoch duration was not an exact multiple of 5 s, the final window was aligned with the end of the block. Each window was assigned the label corresponding to its FP generator. The resulting dataset therefore consisted of 5000-sample signal segments (5 s at 1 kHz sampling rate) and their associated FP generator labels.

**Figure 1.**
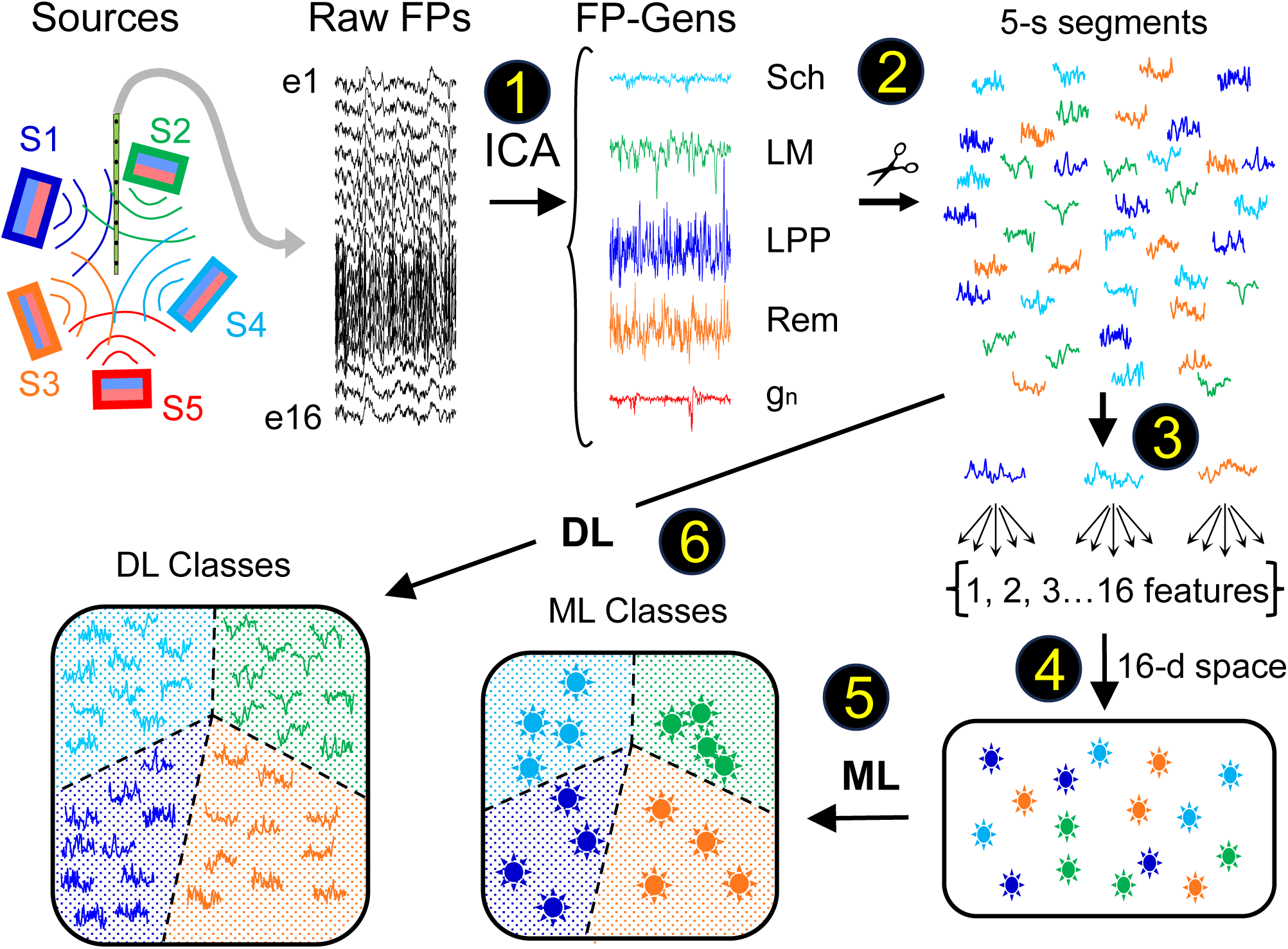
Analysis pipeline. (1) Raw field potential (raw FPs) recordings (black traces) reflect volume conduction and mixing of potentials generated by multiple sources (schematics on the left). Linear arrays of raw FP recordings were decomposed by independent component analysis (ICA) into FP generators. (2) Selected FP generators were segmented into consecutive 5-s windows. (3) In the feature-based analysis path, each segment was characterized by 16 different metrics. (4) The resulting 16-dimensional feature vectors from all generators and animals were pooled to construct a common feature space. (5) Different machine learning (ML) models were trained to classify these feature-based representations according to their FP generator of origin. (6) In the parallel deep learning (DL) path, raw FP-generator segments were directly fed into end-to-end models for the same classification task.

#### 2.2.2. Dataset partitioning

The main dataset was divided into training and test sets using an 80:20 proportion. This division was performed at the subject level to prevent data leakage between the training and test sets. The partition was performed separately within each dataset subset (Table1), and the resulting training and test partitions were subsequently pooled. Recording duration was also considered to avoid test sets containing disproportionately short recordings. Because an exact 80:20 division was not always possible given the number of animals and their recording durations, the partition was obtained by random sampling until the target proportion was achieved within a tolerance of 5%.

### 2.3. Feature-based models

#### 2.3.1. Feature selection

Feature-based models operated on a predefined set of signal descriptors calculated from each 5-s segment. Candidate features included statistical, spectral, and signal-complexity measures. The initial candidate set was informed by previous work on the characterization of FP generators (Makarov et al., 2023; Muñoz-Arnaiz et al., 2025; Muela et al., 2026), which identified informative features for generator discrimination. Additional descriptors were considered to broaden the representation of signal properties. A screening procedure was then performed to obtain a compact feature set, taking into account both redundancy among candidate descriptors and their contribution to classification performance. This procedure included a preliminary assessment on the training set to evaluate classification performance and avoid potential overfitting associated with increasing feature-set size. Based on this assessment, a common set of 16 features was retained and subsequently used across all feature-based models, ensuring that differences in classification performance did not arise from model-specific feature selection.

The selected features were: signal skewness; the mean, variance, kurtosis, and skewness of the spectrum; the mean of the power spectral density; negentropy; spectral entropy; singular-value-decomposition entropy; permutation entropy; approximate entropy; sample entropy; wavelet entropy; Rényi wavelet entropy; Katz dimension and Higuchi dimension. The formulas can be found in the Supplementary Methods.

#### 2.3.2. Conventional machine-learning classifiers

We evaluated six conventional ML classifiers: Bagging Classifier (BAG), Light Gradient Boosting Machine (LGBM), Quadratic Discriminant Analysis (QDA), Random Forest (RF), Support Vector Machine (SVM), and Extreme Gradient Boosting (XGB). For SVM, BAG, QDA, MLP, RF, we used *Scikit-learn* in Python (Pedregosa et al., 2011). For XGB and LGBM, we used the *xgboost* (Chen et al., 2025) and *lightgbm* (Shi et al., 2025) Python packages, respectively.

#### 2.3.3. Multilayer perceptron

We also evaluated fully connected multilayer perceptrons (MLPs: Rumelhart et al., 1986; Bishop, 2006) using the same 16-feature input vectors as the conventional ML classifiers. Architectures with zero to four hidden layers and 16, 32, or 64 units per layer were examined. All hidden layers within a given architecture had the same width. The configuration without hidden layers provided a linear baseline, whereas the remaining configurations included progressively deeper nonlinear transformations.

The MLPs were implemented and trained using the same general optimization and evaluation framework as the other DL models.

### 2.4. End-to-end deep-learning models

End-to-end models were trained directly on the preprocessed 5-s generator time series, without prior feature extraction.

### 2.41. Convolutional neural networks

We evaluated one-dimensional convolutional neural networks (CNNs) operating along the temporal axis, which impose locality, temporal weight sharing, translation equivariance, and hierarchical composition (Lecun et al., 1998; Kiranyaz et al., 2021). Architectures with three, six, and nine convolutional layers were examined. For each network depth, we considered two kernel-size configurations to investigate different temporal receptive fields, while progressively increasing the number of feature channels. Each convolutional block consisted of a convolution followed by batch normalization, an exponential linear unit activation (ELU), and average pooling. We also examined two classification heads. In the first configuration, we connected the flattened convolutional features directly to the output layer, without an intermediate hidden layer. The second consisted of a fully connected layer with 256 hidden units, followed by an ELU activation and dropout, before the final classification layer. Detailed architectural specifications are provided in Supplementary Table 2.

#### 2.4.2 Transformers

We evaluated Transformer encoder architectures operating on patch-based representations of the input time series (Nie et al., 2017). Unlike CNNs, Transformers impose weaker locality biases, leveraging self-attention to capture global dependencies across temporal patches while incorporating positional encodings to preserve temporal ordering (Vaswani et al., 2017; Dosovitskiy et al., 2020). Each 5-s signal was divided into patches of 250 samples, considering both non-overlapping and 50%-overlapping configurations. The patches were linearly projected into a latent embedding space, and a learnable classification token was prepended to the sequence. Fixed sinusoidal positional encodings were then added to incorporate positional information.

We examined architectures comprising two or four Transformer encoder layers, varying the embedding dimension and the number of attention heads while maintaining a fixed dimension per head. Two alternative classification heads were evaluated using the final representation of the classification token. The first mapped this representation directly to the output classes through a fully connected layer, without intermediate hidden layers. The second incorporated an additional fully connected layer with 256 hidden units, followed by GELU activation and dropout before the output layer. Detailed architectural specifications are provided in Supplementary Table 2.

### 2.5. Model training and evaluation

#### 2.5.1 Hyperparameter optimization

Hyperparameters were optimized on the training set using leave-one-subject-out cross-validation, with each validation fold comprising all segments from one held-out subject to prevent data leakage (Del Pup et al., 2025). The best configuration for each model was selected based on mean validation accuracy across folds.

For classical machine-learning methods, we used Optuna (Akuna et al., 2019), with search spaces detailed in the Supplementary Material. For deep-learning models, we prioritized architecture size to mitigate overfitting, progressively increasing model capacity until validation performance no longer improved.

#### 2.5.2 Deep-learning training

Deep-learning models were trained using a batch size of 64 for a maximum of 50 epochs. Early stopping was applied with a patience of 20 epochs. The learning rate was controlled using a cosine annealing scheduler with an initial warm-up phase of 10 epochs. Initial learning rates were set to 1e-2, 1e-5, and 1e-3 for MLPs, CNNs, and Transformers, respectively. The minimum learning rate was 1e-8 for all model families.

#### 2.5.3 Final model evaluation

Following hyperparameter optimization, the selected configuration of each model was retrained using the entire training set. Final performance was assessed once on the independent test set, which had not been used during training or hyperparameter optimization.

### 2.6 Statiscs

Differences in classification accuracy among the six feature-based models were assessed using the Friedman test, treating each animal as an independent experimental unit. Kendall’s W was calculated as an effect-size measure. When significant overall differences were detected, pairwise comparisons were performed using Wilcoxon signed-rank tests, with Holm correction for multiple comparisons.

Spearman’s rank correlation was used to examine the relationship between model complexity and classification accuracy across animals. Correlations were calculated separately for each animal to assess whether the association between model complexity and classification performance was consistent across subjects.

To account for the hierarchical structure of the classification data and the dependence arising from evaluating all models on the same segments, differences in classification performance were additionally assessed using a generalized linear mixed-effects model (GLMM). The binary classification outcome (correct or incorrect prediction) was modelled using a binomial distribution and a logit link. Model identity and ground-truth class were included as fixed effects. Random intercepts were included for recording and segment identity, as well as for rat-by-model and recording-by-model combinations, to account for the dependence among observations and allow model performance to vary across animals and recordings. Pairwise comparisons between models were adjusted for multiple testing using the Holm method. Model fitting and diagnostics were performed using the *glmmTMB* package in R. Statistical significance was set at p < 0.05 (Brooks et al., 2017; McGillycuddy et al., 2025).

## 3. RESULTS

The analysis pipeline is summarized in Figure 1. FP generators were obtained from linear hippocampal recordings along the CA1/DG axis, and those showing greater stability over time and across animals were selected for analysis. Long recording epochs were then segmented into 5-s segments, which followed two parallel analysis paths. In one path, each segment was characterized using a battery of metrics (up to 16, see Methods), yielding a multidimensional feature space that provided a set of complementary descriptors of FP-generator activities. These feature-based representations were subsequently used as input to machine learning (ML) models for classification. In the other path, the raw 5-s segments were directly fed into deep learning (DL) models.

### 3.1. Classification performance

The classification performance of all models is summarized in Figure 2. Overall, raw-signal models outperformed feature-based models. TRF achieved the highest accuracy (94.21% ± 2.75%), closely followed by CNN (93.85% ± 4.45%), with very similar accuracy. Among feature-based models, accuracy ranged from 80.38% ± 11.86% for QDA to 85.83% ± 9.75% for LGBM. Thus, the best-performing raw-signal model exceeded the best feature-based model by 8.38 percentage points. Feature-based models also showed greater variability in classification performance across animals.

**Figure 2.**
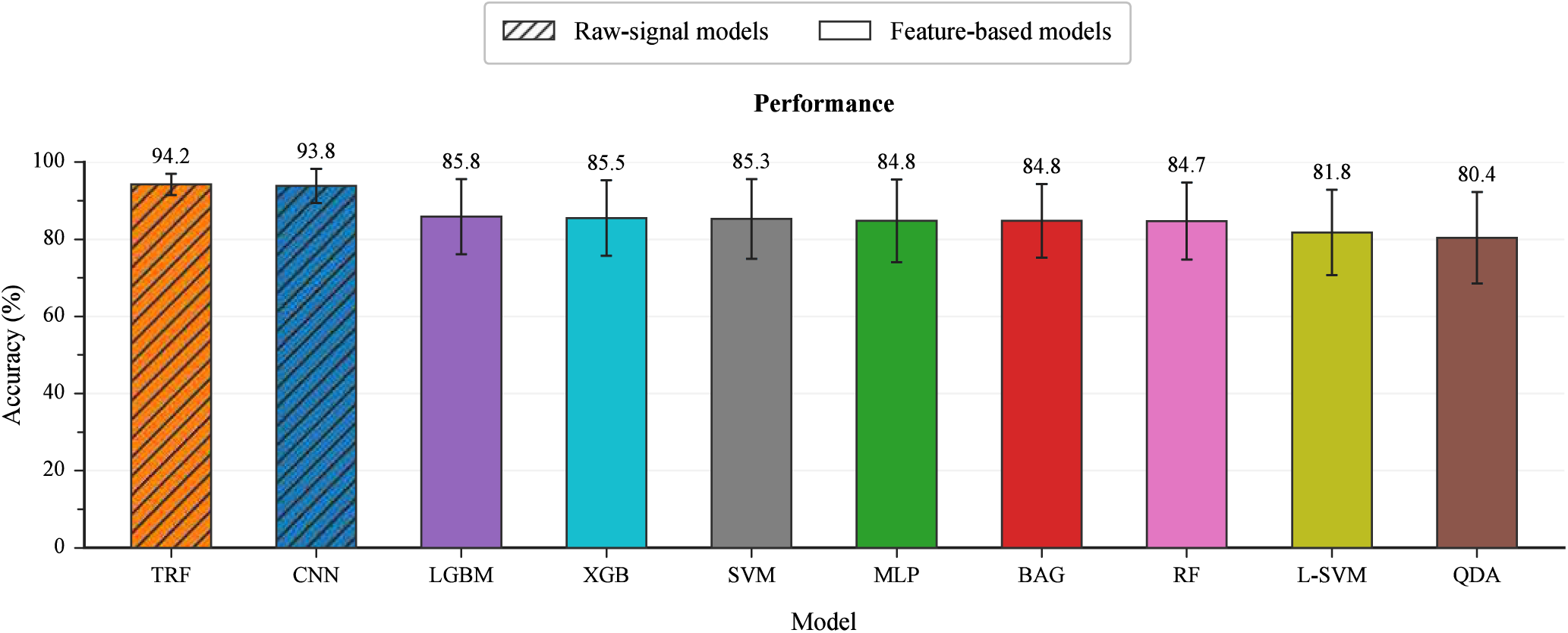
Performance comparison of the evaluated models. The bar chart shows the mean rat-level accuracy achieved by each model, with models ranked in descending order of mean accuracy. Accuracy is computed independently for each rat and subsequently averaged across all rats in the test set. Error bars indicate one standard deviation across rats, and numerical values above the bars denote the corresponding mean accuracies. Hatched bars represent raw-signal models, whereas non-hatched bars represent feature-based models.

Figure 3 shows the confusion matrices of the two best-performing raw-signal models, TRF and CNN, and feature-based models, LGBM and XGB. LPP and Remote generators were the most readily classified, with recall values above 97% across all four models. Most classification errors involved SCH and LM generators. The raw-signal models performed better on these two classes, with recall values of 94.18% (SCH) and 83.83% (LM) for TRF, and 96.43 (SCH) and 80.63% (LM) for CNN. The main confusion occurred between these two classes, with 10.12% of LM samples classified as SCH by TRF and 15.95% by CNN. The feature-based models showed lower recall for both classes. LGBM archived 82.65% for SCH and 64.69% for LM, while XGB obtained 81.36% and 64.28%, respectively. Both models showed similar errors, with 16.00% and 17.57% of LM samples classified as SCH, 15.96% and 14.58% as Remote, and 10.01% and 11.49% of SCH samples classified as LPP.

**Figure 3.**
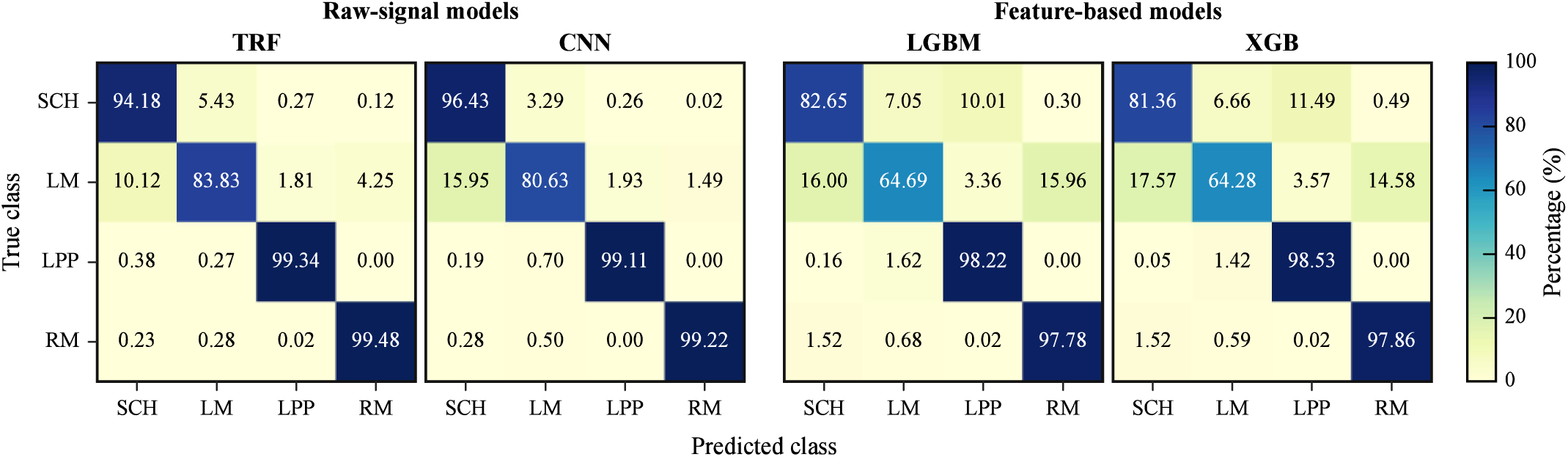
Mean normalized confusion matrices of the best-performing models. The two confusion matrices on the left correspond to TRF and CNN, the two best-performing raw-signal models, whereas those on the right correspond to LGBM and XGB, the two best-performing feature-based models. For each model, a row-normalized confusion matrix is computed independently for each rat and subsequently averaged across all rats in the test set, ensuring equal contribution from each rat. Rows represent the true classes, and columns represent the predicted classes. Values along the main diagonal indicate the mean percentage of correct classifications for each class, whereas off-diagonal values represent the corresponding misclassification percentages. All matrices share a common color scale ranging from 0 to 100%.

To further characterize class discrimination, Figure 4 shows the ROC curves for all models. For the feature-based models, SCH and LM generally showed lower discriminative performance than LPP and Remote.

**Figure 4.**
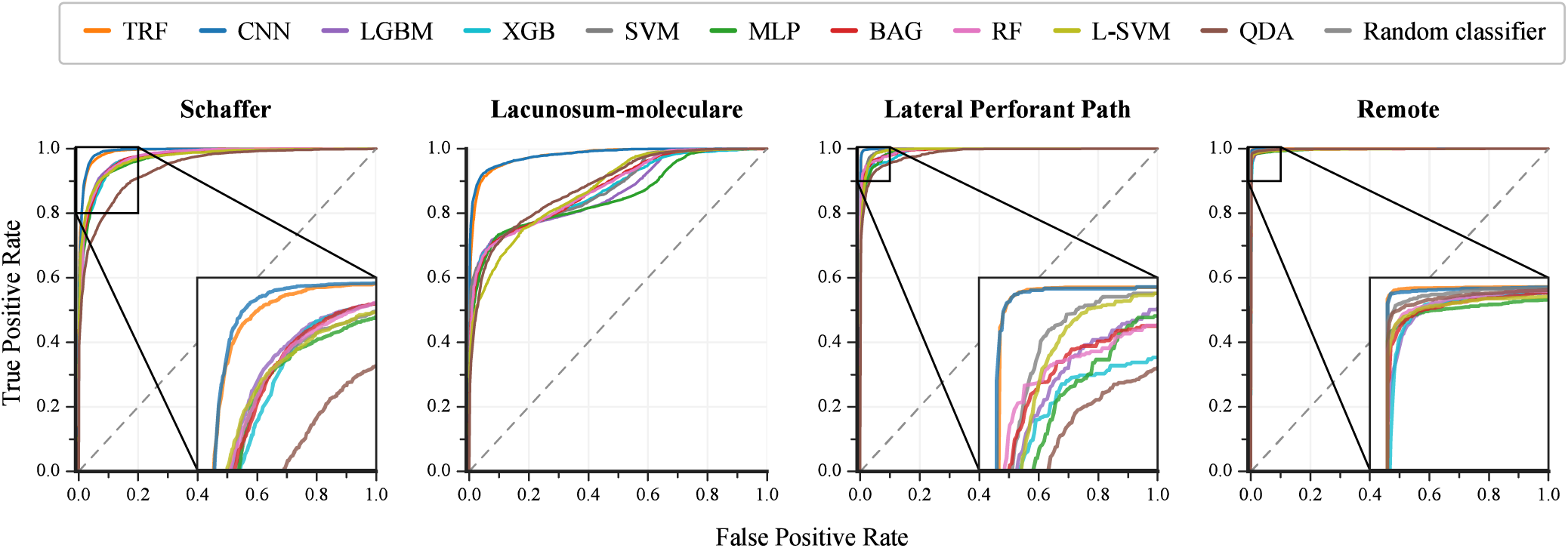
Mean ROC curves of the evaluated models for each class. The four panels correspond to the classes considered: Schaffer, Lacunosum-moleculare, Lateral Perforant Path, and Remote. For each model and class, a one-vs-rest ROC curve is computed independently for each rat. The resulting curves are interpolated onto a common false positive rate grid, and the corresponding true positive rates are averaged across all rats in the test set. Each curve therefore represents the mean rat-level relationship between the true positive rate and the false positive rate. The dashed diagonal line represents the expected performance of a random classifier, whereas the insets provide magnified views of regions with low false positive rates and high true positive rates to facilitate comparison between models.

The raw-signal models showed consistently high discriminative performance across classes, with ROC curves generally close to the upper-left corner. Their similar profiles were consistent with the comparable classification accuracies obtained by CNN and TRF.

### 3.2. Statistical comparison of model performance

The overall comparison of the six feature-based models revealed significant differences in classification accuracy (Friedman test, χ² = 16.65, p = 0.0106; Kendall’s W = 0.463). However, none of the pairwise comparisons remained significant after Holm correction. The relationship between model complexity and accuracy also varied across subjects, as indicated by the heterogeneous Spearman correlations (ρ ranging from −0.09 to 0.77). Only one subject showed a nominally significant positive correlation (ρ = 0.775, p = 0.041), providing no consistent evidence of an association between model complexity and classification accuracy across subjects.

Differences in classification performance between models were further assessed using a generalized linear mixed-effects model (GLMM) with a binomial distribution and a logit link. This approach accounted for the dependence among observations arising from the evaluation of all models on the same segments, which were hierarchically organized within recordings. The fixed effects were classification model, ground-truth class, and their interaction. Random intercepts were included for segment, recording, and each recording-by-model combination. Treating these observations as statistically independent could otherwise lead to pseudoreplication (Lazic, 2010).

The model converged, and the Hessian matrix was positive definite. No singularity was detected in the random-effects structure. The dispersion analysis indicated statistically significant overdispersion (dispersion ratio = 1.010, p < 0.001), although the deviation from the expected binomial dispersion was small (approximately 1%).

Averaged equally across the four ground-truth classes, TRF and CNN had the highest GLMM-estimated accuracies (99.41% and 98.66%, respectively). Among the eight feature-based models, estimated accuracy ranged from 86.50% for L-SVM to 95.51% for MLP. Mean estimated accuracy was 8.21 percentage points higher for the two raw-signal models than for the eight feature-based models (95% CI 0.48 to 15.94 percentage points, p = 0.0374). None of the 17 exploratory pairwise comparisons reached statistical significance after Holm correction, including the comparison between CNN and TRF (Holm-adjusted p = 0.832).

For TRF, the log-odds of correct classification were lower for LM (β = −9.08) and Sch (β = −3.82) than for LPP (both p < 0.001). The difference between Remote and LPP for TRF was not statistically significant (β = 0.33, p = 0.451).

### 3.3. Confidence-based analysis of classification errors

The performance differences observed between raw-signal and feature-based models raise the question of whether segments misclassified by feature-based models are also difficult to classify using raw-signal models. To address this question, we examined the confidence of TRF predictions for segments incorrectly classified by the best-performing feature-based model.

As a preliminary step, we evaluated the calibration of TRF using the Expected Calibration Error (ECE) and the Brier Score (BS). TRF achieved an ECE of 0.0266 and a BS of 0.1038.

We then analyzed the errors made by LGBM, the best-performing feature-based model. For each segment incorrectly classified by LGBM, we assessed whether TRF classified it correctly and examined the confidence associated with its prediction. Figure 5 shows the distribution of these errors according to TRF confidence, distinguishing between segments correctly classified by TRF and those misclassified by both models.

**Figure 5.**
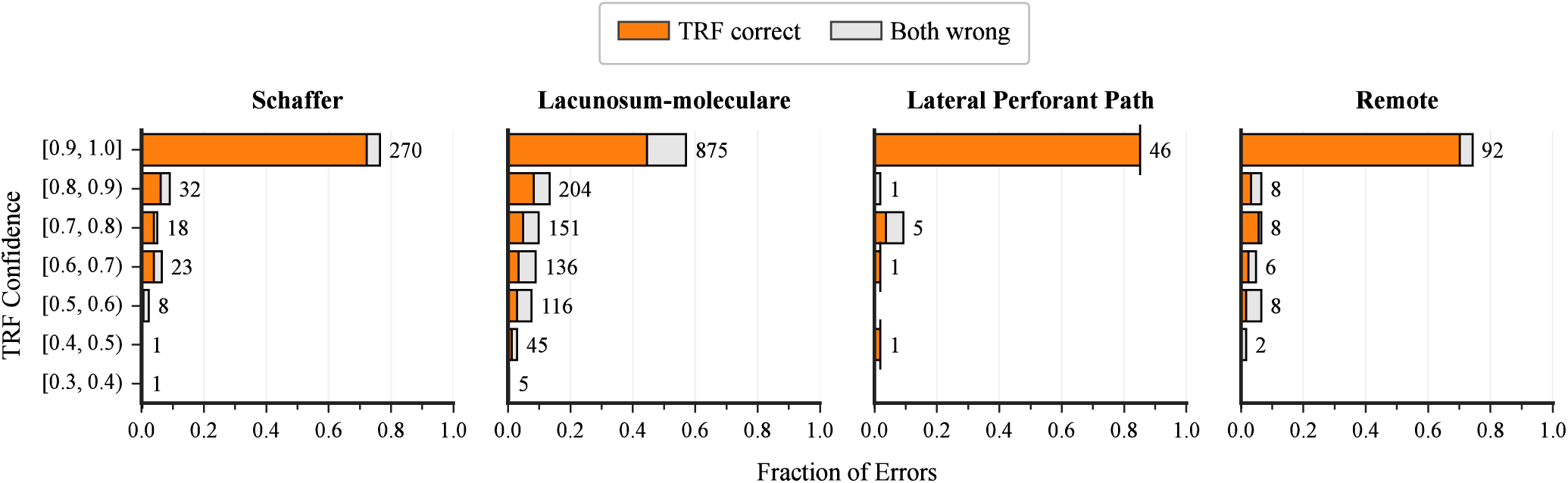
Confidence-based analysis of LGBM misclassifications. For each class, only samples misclassified by LGBM are considered and grouped into confidence intervals according to the maximum class probability predicted by TRF. Orange segments represent samples correctly classified by TRF, whereas grey segments represent samples misclassified by both models. The length of each segment indicates the fraction of the total LGBM misclassifications within the corresponding true class. Numerical values alongside the bars indicate the absolute number of samples in each confidence interval. The four panels allow for a class-wise comparison of the ability of TRF to correctly classify samples misclassified by LGBM at different confidence levels.

TRF correctly classified 71.01% of the segments misclassified by LGBM. Moreover, 51.91% of all LGBM errors were correctly classified by TRF with confidence between 0.9 and 1.0. This proportion increased to 59.33% when considering predictions with confidence equal to or greater than 0.8.

These results indicate that a substantial proportion of the segments misclassified by the feature-based model could nevertheless be correctly classified by TRF, often with high confidence. Thus, errors made by LGBM do not necessarily correspond to segments that are difficult to classify across models, but may instead reflect limitations in the information captured by the feature-based representation.

## 4 DISCUSSION

The predominant irregular temporal activity in FPs contains sufficient information in short signal segments to discriminating among their component FP generators mixed in the volume. Predefined features captured a substantial part of this information, although raw-signal models achieved better classification performance, indicating that additional discriminative information can be extracted directly from the signals. The performance differences also depended on the classification algorithm and the FP generator considered, while the ability to discriminate among them was maintained across animals recorded by different experimenters and under diverse experimental conditions.

### 4.1. Regional discriminability of irregular activity and generalization across animals

Several factors may contribute to the irregular appearance of FPs and EEG signals, including destructive interference among potentials generated by co-activated sources within the recording volume (Ahlfors et al., 2010; Herreras, 2016), whose relative contributions vary with recording location (Herreras et al., 2022, 2023). Irregularity may also reflect the sparse and stochastic nature of neural coding and the integration of diverse synaptic, cellular, and network properties across spatial scales (Muñoz-Arnaiz et al., 2026). Understanding these contributions remains challenging, as conventional spectral features may provide limited insight into temporal organization, while less conventional descriptors are not always readily interpretable in physiological terms (Thivierge and Cisek, 2008; He et al., 2010).

The separation of FP generators can help reveal signal properties that may be obscured by spatial mixing. For example, we previously showed that changes in correlation dimension (one of the features included in the present study) preceding epileptic seizures could not be consistently predicted from raw FPs, whereas their direction became predictable when analyzing separated FP generators (Makarov et al., 2023). Previous machine-learning studies further showed that generator identity is reflected in characteristic signal patterns, including stable properties and changes induced by exogenous behavioral manipulations (Muela et al., 2026). The present results extend these findings by demonstrating that, despite their irregular temporal organization and apparent similarity in short segments, generator-specific information can be captured by different models using predefined signal descriptors.

An important aspect of this study is that each model was evaluated on animals not included during training. Moreover, the dataset encompassed substantial experimental heterogeneity, including recordings obtained by four different experimenters over a period of 10 years, across different experimental series, using different acquisition systems and electrode types. The ability to discriminate among FP generators in held-out animals indicates that the relevant signal patterns are not restricted to individual subjects or to a single experimental configuration. Although the individual contribution of each experimental factor was not assessed, the consistently high discrimination performance across models indicates that generator-related information remained detectable across the range of experimental conditions represented in the dataset. Thus, these results support generalization across animals recorded under diverse experimental conditions, although they do not establish independence from any particular experimental factor.

A note of caution is warranted regarding anesthetic dosage. In the main dataset, anesthetic dosage was kept relatively consistent across experimental series. This is important because both anesthetic depth and wakefulness are known to influence the overall electrographic pattern (Kuo and Leung, 2017). In addition, changes in the composition of the underlying sources may affect ICA performance (Makarova et al., 2011). We therefore excluded recordings obtained under substantially different state conditions from the main analysis, since any resulting differences in classification performance could not be readily attributed to the ICA decomposition, feature selection, or the models’ ability to discriminate among activity patterns.

In a preliminary test using a group of four rats recorded in another laboratory (V. López-Madrona and S. Canals) under a higher anesthetic dose (2 g/kg), all generators except the Schaffer generator were correctly identified by all models, whereas the TRF correctly identified all four generators. Although this observation is too limited to support firm conclusions, it suggests that model generalization may be differentially affected by changes in experimental conditions, potentially depending on the generator being classified. In particular, it raises the possibility that some models may be more robust than others to variations in anesthetic state. Assessing this possibility will require a substantially larger dataset in which different anesthetic conditions are systematically represented.

### 4.2. Discriminative information captured by predefined features and raw-signal models

A question addressed in this study was how much generator-specific information is captured by predefined signal features. The feature-based models achieved substantial classification performance, demonstrating that the selected temporal, spectral, and nonlinear properties contain relevant information for distinguishing among generators. However, the range of potentially informative signal features is broad, and different signals or activity regimes may be better represented by different combinations of features, depending, for instance, on the generators being compared. Thus, feature selection may need to be tailored to the specific signals and discrimination task.

Nevertheless, raw-signal models generally achieved higher classification performance. Operating directly on the temporal signals, these models can exploit information without requiring its prior reduction to a predefined set of descriptors. Their superior performance therefore suggests that the selected features do not capture all the discriminative information available in the original signals.

The confidence-based analysis provides further support for this interpretation. A substantial proportion of the segments incorrectly classified by LGBM were correctly classified by TRF, often with high prediction confidence. This indicates that many segments misclassified by the feature-based model still contain information that can be exploited by a model operating directly on the signal. Thus, at least part of the performance difference between the two approaches may arise from limitations of the predefined representation, rather than solely from the presence of intrinsically difficult segments. However, the present analysis does not identify which specific signal properties account for the additional discriminative information exploited by TRF.

The similar performance of CNN and TRF raises the possibility that the advantage of raw-signal models derives primarily from their ability to learn directly from temporal data, rather than from a particular architectural design. However, determining whether these architectures rely on similar signal patterns would require further investigation of their internal representations and decision processes.

### 4.3. Implications, limitations and future directions

The present results support the use of computational approaches to identify generator-related information in irregular FP activity (Muñoz et al., 2025; Muela et al., 2026). The ability to discriminate among generators from short segments and across different animals and experimental conditions provides a basis for investigating the spatial origin of electrophysiological signals using their temporal properties, rather than relying exclusively on direct inspection or predefined signal descriptors.

An important future direction is to determine which temporal patterns are responsible for the discrimination achieved by raw-signal models. Although deep-learning approaches can identify complex patterns directly from the data, the physiological meaning of these patterns is not necessarily evident from their classification performance alone. Explainable artificial intelligence and mechanistic interpretability approaches could help identify the signal properties and internal computations supporting model predictions (Arrieta et al., 2020; Samek et al., 2021; Bereska and Gavves, 2024). In particular, examining which temporal features contribute to the distinction between generators could help establish links between computationally identified patterns and the underlying electrophysiological processes.

These approaches may have clinical implications beyond epilepsy. The applicability of FP generators to human intracranial recordings has already been demonstrated (Makarova et al., 2025), providing a basis for extending the present classification framework to clinical data. Their spatial profiles allow local and remote contributions to be distinguished, providing information about the anatomical origin of recorded activity that may be obscured by conventional bipolar recordings, while their temporal fluctuations can be characterized with less degradation than in raw FPs. The present results suggest that combining feature-based ML classifiers, which offer a practical means of exploring and discriminating FP generators, with DL approaches operating directly on generator signals could further improve the characterization of complex activity. One potential application is the discrimination of physiological and pathological activity, even when both exhibit irregular temporal patterns. Rather than relying on irregularity itself as an indicator of pathology, generator-based analysis could help identify differences in the temporal and spectral organization of activity and in the relative contributions of distinct generators. In epilepsy, for example, this approach might help characterize the regions contributing to seizure onset and propagation, although distinguishing generators of pathological activity from those recruited secondarily will require further investigation.

Several limitations should be considered. First, the present study focused on four generators and a specific experimental preparation, and whether the approach can discriminate a broader range of generators, brain regions, or physiological and pathological conditions remains to be established. Second, although the internal dataset included substantial variability across animals and experimental configurations, independent external validation under systematically different recording conditions is still lacking. Third, the physiological properties underlying the additional discriminative information exploited by raw-signal models remain unknown. Future studies should address these limitations by extending the approach to additional recording configurations and pathological models, while investigating the signal patterns and physiological mechanisms underlying model predictions. Combining generator-based classification with interpretable computational analyses may ultimately improve our understanding of the spatial organization of extracellular signals and their use in experimental and clinical neuroscience.

## Acknowledgements

We thank former lab members Nuria Benito, and Marta Montero and Julia Isidro from the CIAL-CSIC who performed some of the experiments from which the FP recordings used in this study were obtained.

## Author Contributions

Conceptualization: MGL, OH; Model construction: NCT, ARF, RAM, JBA; Investigation: NCT, AFR, RAM, FA, JM; Funding acquisition: MGL, OH; Writing original draft NCT, OH; Writing-review & editing: NCT, AFR, RAM, JBA, FA, JM, MGL, OH.

---

**Supplementary Table 1.** Description of variables across experimental series. Experimenter: FA (Faezeh Ashouri), NB (Nuria Benito), MNA (Marta Montero-Atalaya), and JI (Julia Isidro). Sex: F (female). Age is given in months. Anesthesia dose is expressed in g/kg body weight of urethane. Left/right indicates the recorded hemisphere. Probe: N (Neuronexus, Ann Arbor, Michigan, USA) and A (Atlas Neuro, Leuven, Belgium). Channel count indicates the number of contacts (16 or 32), arranged linearly; contact spacing is given in μm. Hardware: OE (Open Ephys, Lisbon, Portugal) and MCS (Multi Channel Systems, Reutlingen, Germany). Sampling rate is given in Hz. Segments indicates the number of 5-s segments into which each recording epoch was divided for analysis. Time refers to recording duration in minutes. Dataset identifies the combinations of experimental variables specified in Table 1 of the main text.

| Experimenter | Exp. Name | Sex | Age | Anest | L/R | Probe | Ch count/spacing | shanks | Hardw | SR | Segments | Rec time | Dataset |
|---|---|---|---|---|---|---|---|---|---|---|---|---|---|
| FA | 06.08.2025- 1h control- downsampled- 33 64 cured_ICA-1 | F | 3 | 1.5 | R | N | 16/100 | Twin | OE | 2000 | 2560 | 213 | D |
| FA | 09.09.2025-1h-control-downsampled- 1 32 cured_ICA-1 | F | 3 | 1.5 | L | N | 16/100 | Twin | OE | 2000 | 2316 | 193 | D |
| FA | 09.09.2025-1h-control-downsampled- 33-64_ ICA-1 | F | 3 | 1.5 | R | N | 16/100 | Twin | OE | 2000 | 2304 | 192 | D |
| FA | 14.07.2025 1 h control_33-64_channels_SinTheta_ICA-1 | F | 3 | 1.5 | R | N | 16/100 | Twin | OE | 2000 | 1796 | 150 | D |
| FA | 16.09.25-1h-control- 1 32 channels_downsampled_ICA-1 | F | 3 | 1.5 | R | N | 16/100 | Twin | OE | 2000 | 2380 | 199 | D |
| FA | 16.09.25-1h-control- 33 64 channels_downsampled_ICA-1 | F | 3 | 1.5 | R | N | 16/100 | Twin | OE | 2000 | 2324 | 194 | D |
| FA | 19.08.2025-control-1-32-SinTheta_ICA-1 | F | 3 | 1.5 | L | N | 16/100 | Twin | OE | 2000 | 888 | 74 | D |
| FA | 19.08.2025-control-33-64-SinTheta_ICA-1 | F | 3 | 1.5 | R | N | 16/100 | Twin | OE | 2000 | 888 | 74 | D |
| FA | 24.09.2025-1h control (50-60 excluded)- 33 64- downsampled_ICA-1 | F | 3 | 1.5 | R | N | 16/100 | Twin | OE | 2000 | 1416 | 118 | D |
| FA | 24.09.2025-1h-control (50-60 excluded)- 1 32 cured-downsampled_I | F | 3 | 1.5 | L | N | 16/100 | Twin | OE | 2000 | 1072 | 89 | D |
| FA | 25.09.2025-50 mins control-33 64 channels- downsampled_ICA-1 | F | 3 | 1.5 | R | N | 16/100 | Twin | OE | 2000 | 1012 | 84 | D |
| FA | 29.08.2025- 1h control_1-32_SINTHETA_ICA-1 | F | 3 | 1.5 | L | N | 16/100 | Twin | OE | 2000 | 2172 | 181 | D |
| FA | 29.08.2025- 1h control_33-64_SINTHETA_ICA-1 | F | 3 | 1.5 | R | N | 16/100 | Twin | OE | 2000 | 1992 | 166 | D |
| NB | Rata109_4Lancetas_TodosGnrs.mat | F | 3 | 1.2 | L | N | 16/50 | Twin | MCS | 4000 | 48 | 4 | C |
| NB | Rata111_4Lancetas_TodosGnrs.mat | F | 3 | 1.2 | L | N | 16/50 | Twin | MCS | 4000 | 48 | 4 | C |
| NB | Rata121_4Lancetas_TodosGnrs.mat | F | 3 | 1.2 | L | N | 16/50 | Twin | MCS | 2000 | 56 | 4.7 | C |
| NB | Rata122_2Lancetas_TodosGnrs.mat (solo 2 lancetas) | F | 3 | 1.2 | L | N | 16/50 | Twin | MCS | 4000 | 48 | 4 | C |
| NB | Rata124_4Lancetas_TodosGnrs.mat | F | 3 | 1.2 | L | N | 16/50 | Twin | MCS | 4000 | 48 | 4 | C |
| FA | 07.07.2025-control-only16chs-SinTheta_ICA-1 | F | 3 | 1.5 | R | N | 16/100 | Twin | OE | 2000 | 1600 | 133 | C |
| FA | 16.07.2025- 1h control- 33-48 channels_ICA-1 | F | 3 | 1.5 | R | N | 16/100 | Twin | OE | 2000 | 2176 | 181 | C |
| FA | 16.07.2025- 1h control- 49-64 channels_ICA-1 | F | 3 | 1.5 | R | N | 16/100 | Twin | OE | 2000 | 1792 | 149 | C |
| MMA | Raton_3_noTheta_ICA-J.mat | F | 4 to 5 | 1.5 | R | A | 29/65 | Simple | MCS | 1000 | 592 | 49 | A |
| MMA | Raton_5_noTheta_ICA-J.mat | F | 4 to 5 | 1.5 | R | A | 29/65 | Simple | MCS | 1000 | 552 | 46 | A |
| MMA | Raton_2_noTheta_ICA-J.mat | F | 4 to 5 | 1.5 | R | A | 30/65 | Simple | MCS | 1000 | 804 | 67 | A |
| MMA | Raton_4_noTheta.mat | F | 4 to 5 | 1.5 | R | A | 30/65 | Simple | MCS | 1000 | 552 | 46 | A |
| JI | Control5-hasta005_noTheta_ThetaMarks_ICA-1.mat | F | 3 | 1.5 | R | N | 31/100 | Simple | OE | 2000 | 3140 | 262 | B |
| JI | Control9_hasta006_noTheta_ThetaMarks_ICA-1.mat | F | 3 | 1.5 | R | N | 31/100 | Simple | OE | 2000 | 3064 | 255 | B |
| MMA | Raton_1_noTheta_ICA-J.mat | F | 4 to 5 | 1.5 | R | A | 32/65 | Simple | OE | 1000 | 1688 | 141 | A |
| MMA | Raton_6_noTheta.mat | F | 4 to 5 | 1.5 | R | A | 32/65 | Simple | MCS | 1000 | 912 | 76 | A |
| JI | Control-0-1-hasta007_noTheta_ThetaMarks_ICA-1.mat | F | 3 | 1.5 | R | N | 32/100 | Simple | MCS | 2000 | 2904 | 242 | B |
| JI | Control-0-2-hasta007_noTheta_ThetaMarks_ICA-1.mat | F | 3 | 1.5 | R | N | 32/100 | Simple | MCS | 2000 | 3516 | 293 | B |
| JI | Control-0-3-hasta010_noTheta_ThetaMarks_ICA-1.mat | F | 3 | 1.5 | R | N | 32/100 | Simple | MCS | 2000 | 1380 | 115 | B |
| JI | Control1_hasta006_noTheta_ThetaMarks_ICA-1.mat | F | 3 | 1.5 | R | N | 32/100 | Simple | MCS | 2000 | 2288 | 191 | B |
| JI | Control10_hasta009_noTheta_ThetaMarks_ICA-1.mat | F | 3 | 1.5 | R | N | 32/100 | Simple | MCS | 2000 | 3600 | 300 | B |
| JI | Control10_masTarde_hasta006_noTheta_ThetaMarks_ICA-1.mat | F | 3 | 1.5 | R | N | 32/100 | Simple | MCS | 2000 | 3276 | 273 | B |
| JI | Control3-hasta007_noTheta_ThetaMarks_ICA-1.mat | F | 3 | 1.5 | R | N | 32/100 | Simple | MCS | 2000 | 1028 | 86 | B |
| JI | Control4-hasta007_noTheta_ThetaMarks_ICA-1.mat | F | 3 | 1.5 | R | N | 32/100 | Simple | MCS | 2000 | 1888 | 157 | B |

**Supplementary Table 2.**
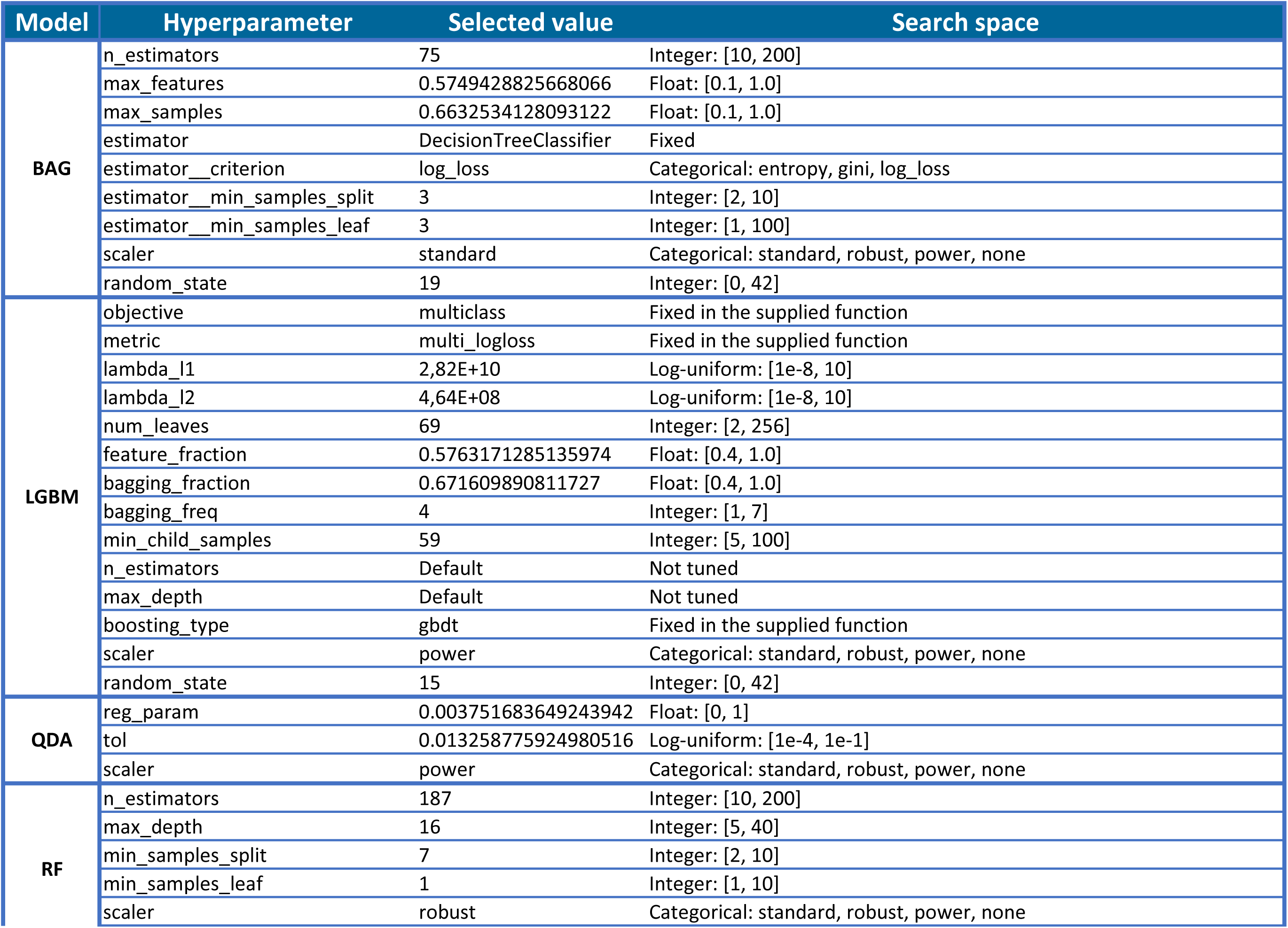
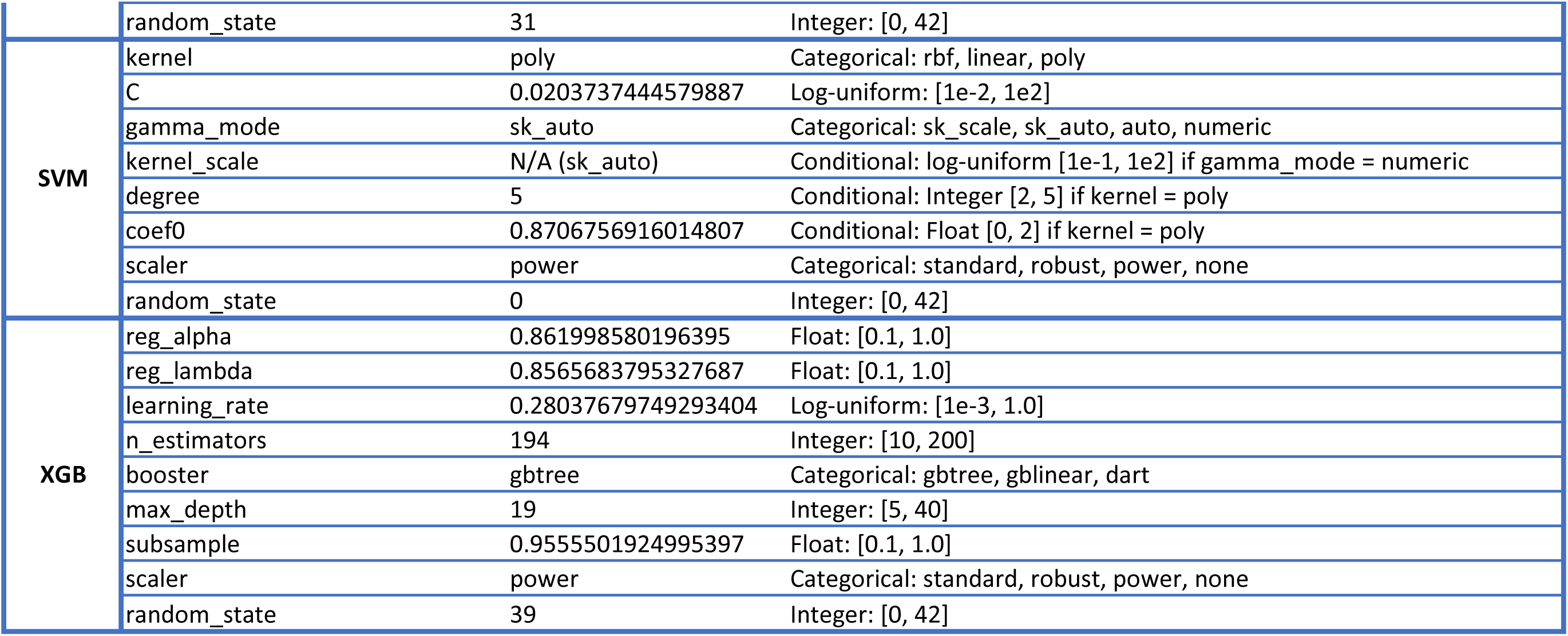
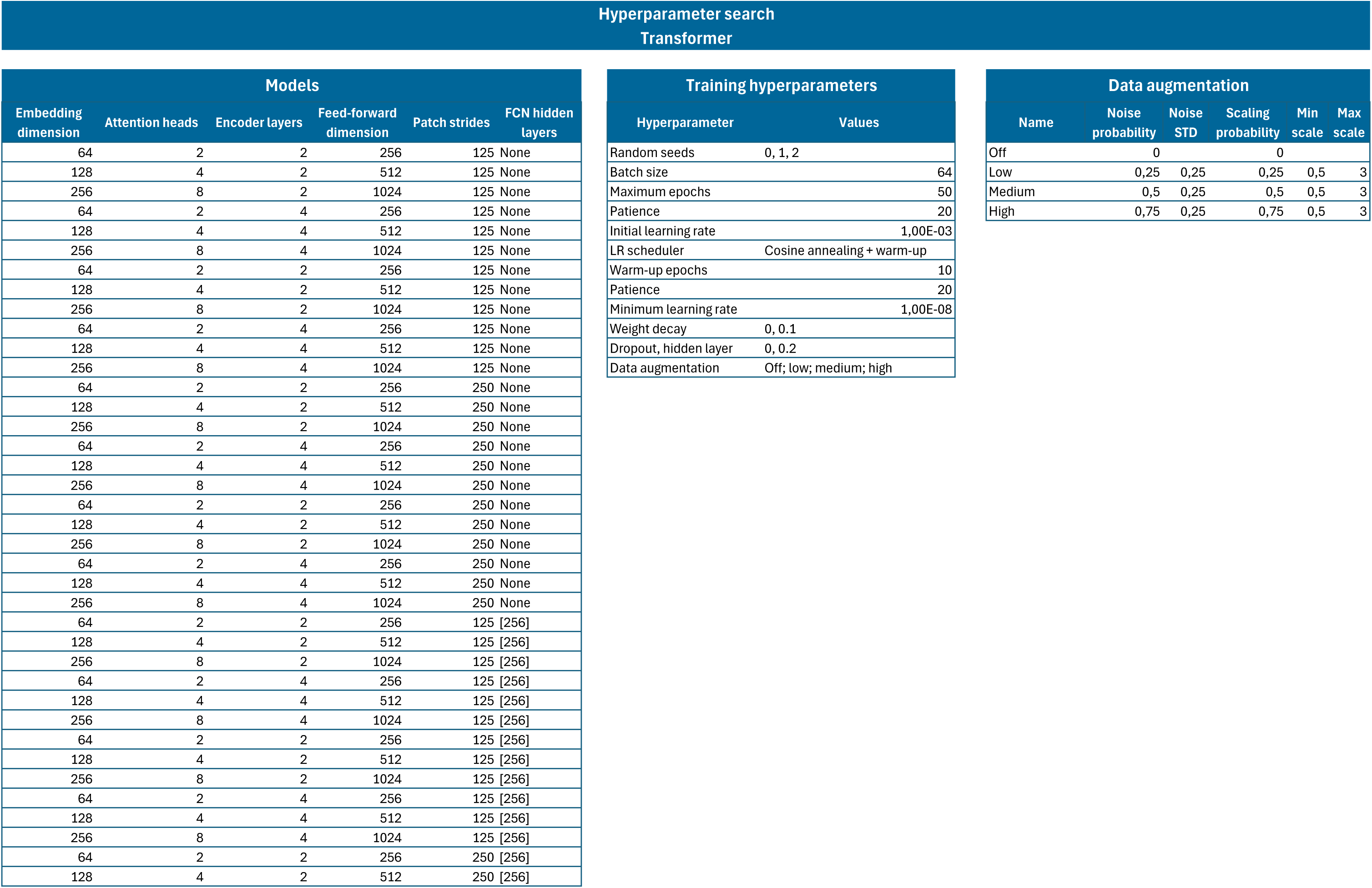
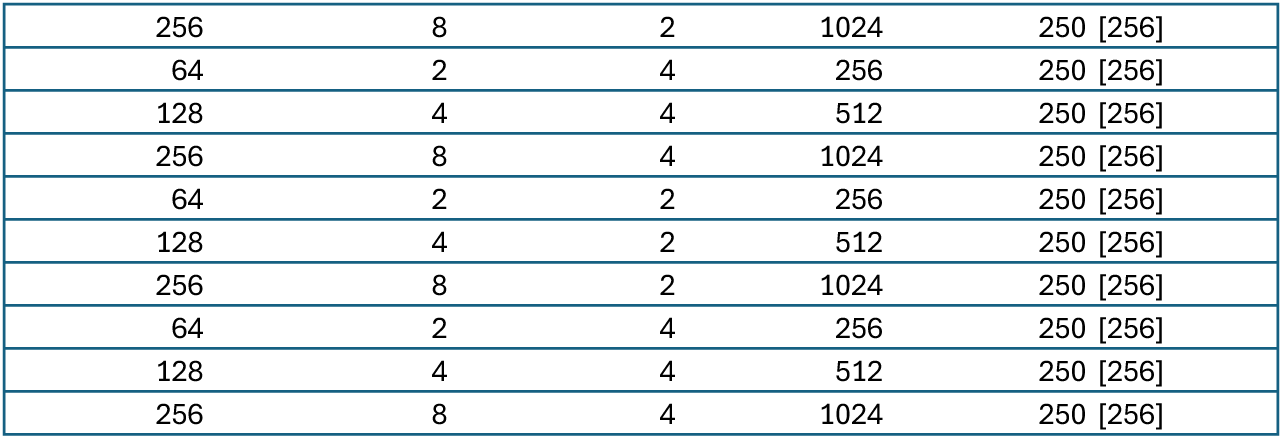
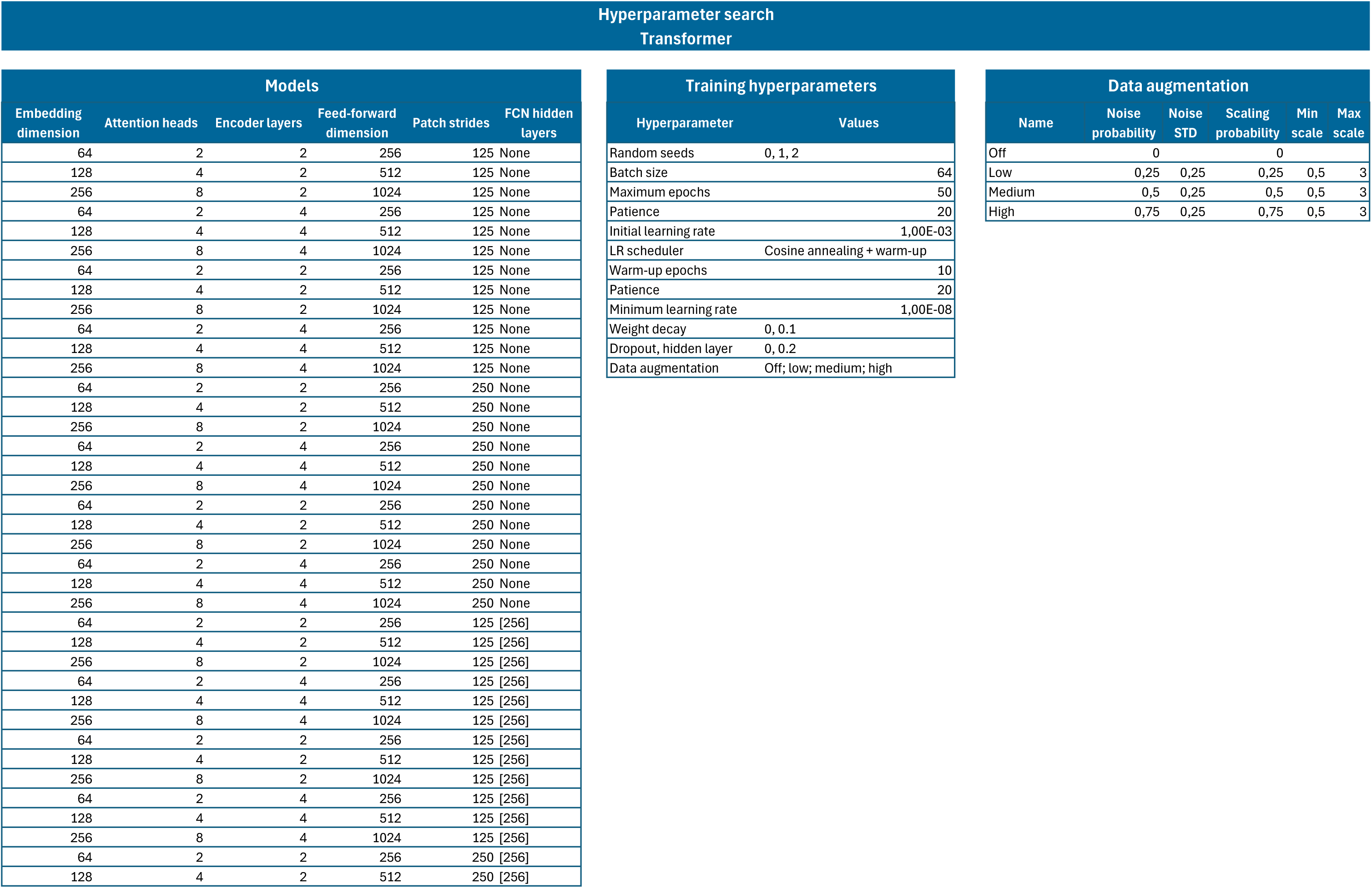
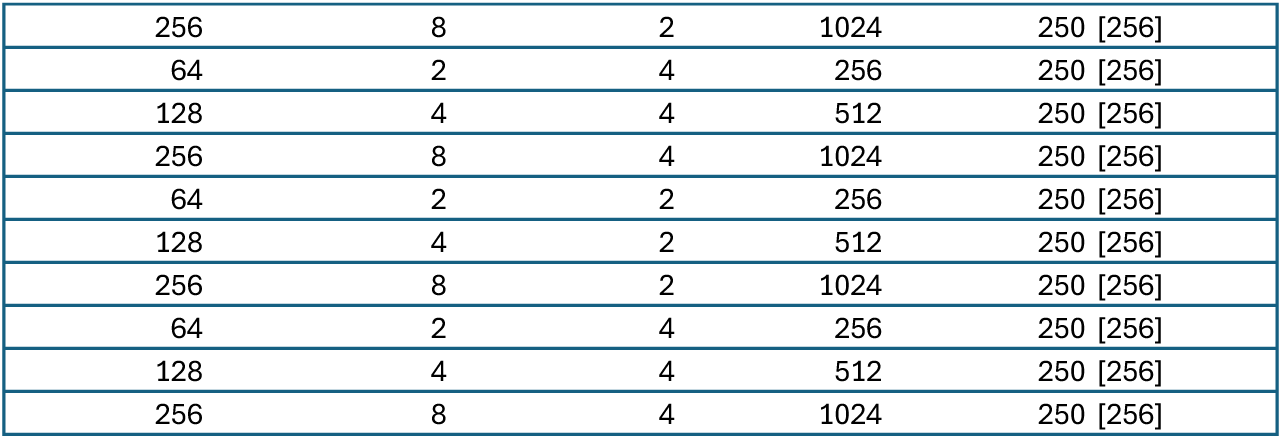
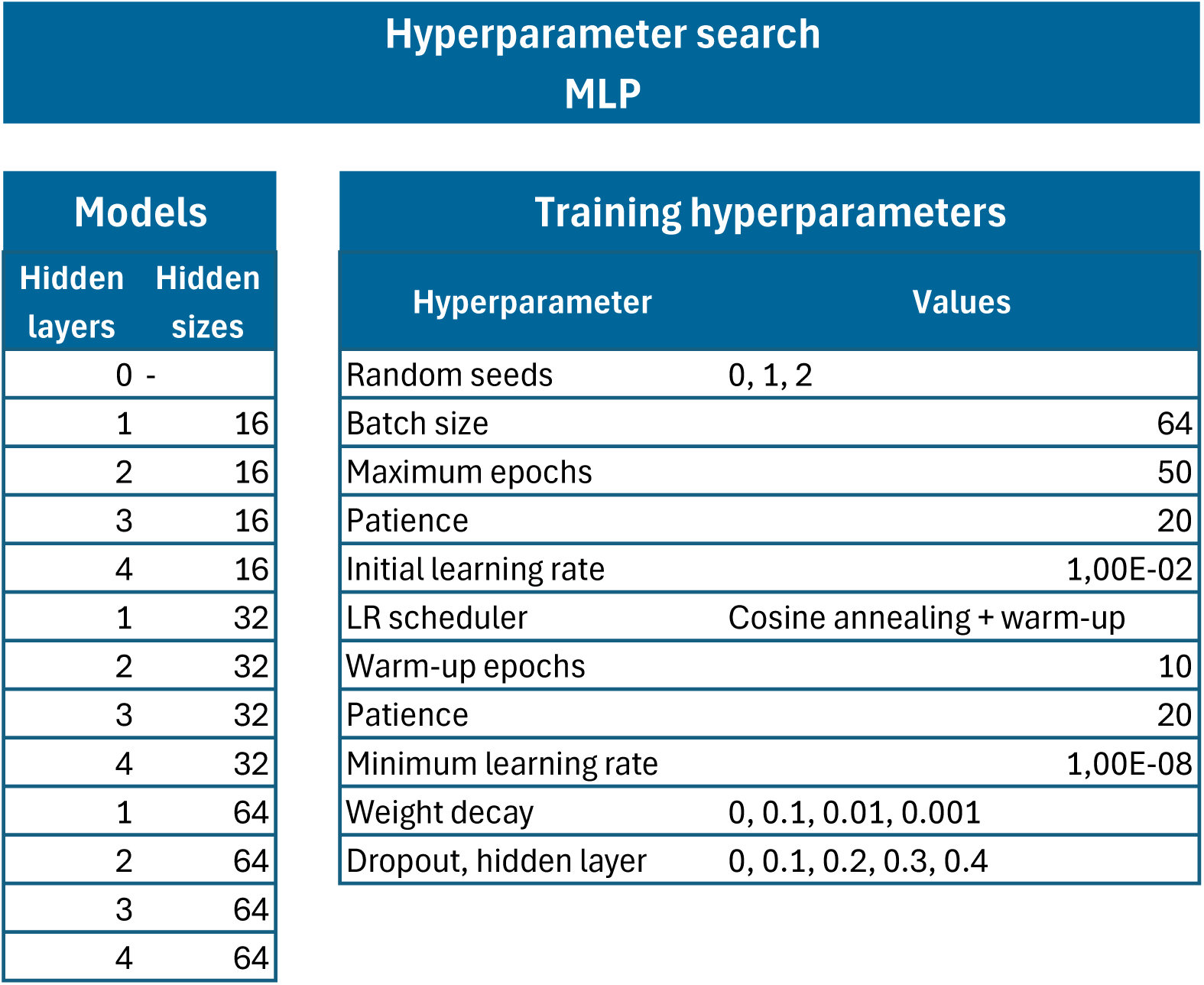
Hyperparameters for ML and DL models.

| Model | Hyperparameter | Selected value | Search space |  |  |
|---|---|---|---|---|---|
| BAG | n_estimators | 75 | Integer: [10, 200] |  |  |
|  | max_features | 0.5749428825668066 | Float: [0.1, 1.0] |  |  |
|  | max_samples | 0.6632534128093122 | Float: [0.1, 1.0] |  |  |
|  | estimator | DecisionTreeClassifier | Fixed |  |  |
|  | estimator__criterion | log_loss | Categorical: entropy, gini, log_loss |  |  |
|  | estimator__min_samples_split | 3 | Integer: [2, 10] |  |  |
|  | estimator__min_samples_leaf | 3 | Integer: [1, 100] |  |  |
|  | scaler | standard | Categorical: standard, robust, power, none |  |  |
|  | random_state | 19 | Integer: [0, 42] |  |  |
| LGBM | objective | multiclass | Fixed in the supplied function |  |  |
|  | metric | multi_logloss | Fixed in the supplied function |  |  |
|  | lambda_l1 | 2,82E+10 | Log-uniform: [1e-8, 10] |  |  |
|  | lambda_l2 | 4,64E+08 | Log-uniform: [1e-8, 10] |  |  |
|  | num_leaves | 69 | Integer: [2, 256] |  |  |
|  | feature_fraction | 0.5763171285135974 | Float: [0.4, 1.0] |  |  |
|  | bagging_fraction | 0.671609890811727 | Float: [0.4, 1.0] |  |  |
|  | bagging_freq | 4 | Integer: [1, 7] |  |  |
|  | min_child_samples | 59 | Integer: [5, 100] |  |  |
|  | n_estimators | Default | Not tuned |  |  |
|  | max_depth | Default | Not tuned |  |  |
|  | boosting_type | gbdt | Fixed in the supplied function |  |  |
|  | scaler | power | Categorical: standard, robust, power, none |  |  |
|  | random_state | 15 | Integer: [0, 42] |  |  |
| QDA | reg_param | 0.003751683649243942 | Float: [0, 1] |  |  |
|  | tol | 0.013258775924980516 | Log-uniform: [1e-4, 1e-1] |  |  |
|  | scaler | power | Categorical: standard, robust, power, none |  |  |
| RF | n_estimators | 187 | Integer: [10, 200] |  |  |
|  | max_depth | 16 | Integer: [5, 40] |  |  |
|  | min_samples_split | 7 | Integer: [2, 10] |  |  |
|  | min_samples_leaf | 1 | Integer: [1, 10] |  |  |
|  | scaler | robust | Categorical: standard, robust, power, none |  |  |
|  | random_state | 31 | Integer: [0, 42] |  |  |
| SVM | kernel | poly | Categorical: rbf, linear, poly |  |  |
|  | C | 0.0203737444579887 | Log-uniform: [1e-2, 1e2] |  |  |
|  | gamma_mode | sk_auto | Categorical: sk_scale, sk_auto, auto, numeric |  |  |
|  | kernel_scale | N/A (sk_auto) | Conditional: log-uniform [1e-1, 1e2] if gamma_mode = numeric |  |  |
|  | degree | 5 | Conditional: Integer [2, 5] if kernel = poly |  |  |
|  | coef0 | 0.8706756916014807 | Conditional: Float [0, 2] if kernel = poly |  |  |
|  | scaler | power | Categorical: standard, robust, power, none |  |  |
|  | random_state | 0 | Integer: [0, 42] |  |  |
| XGB | reg_alpha | 0.861998580196395 | Float: [0.1, 1.0] |  |  |
|  | reg_lambda | 0.8565683795327687 | Float: [0.1, 1.0] |  |  |
|  | learning_rate | 0.28037679749293404 | Log-uniform: [1e-3, 1.0] |  |  |
|  | n_estimators | 194 | Integer: [10, 200] |  |  |
|  | booster | gbtree | Categorical: gbtree, gblinear, dart |  |  |
|  | max_depth | 19 | Integer: [5, 40] |  |  |
|  | subsample | 0.9555501924995397 | Float: [0.1, 1.0] |  |  |
|  | scaler | power | Categorical: standard, robust, power, none |  |  |
|  | random_state | 39 | Integer: [0, 42] |  |  |
| **Hyperparameter search<br>Transformer** |  |  |  |  |  |
| **Models** |  |  |  |  |  |
| **Embedding dimension** | **Attention heads** | **Encoder layers** | **Feed-forward dimension** | **Patch strides** | **FCN hidden layers** |
| 64 | 2 | 2 | 256 | 125 | None |
| 128 | 4 | 2 | 512 | 125 | None |
| 256 | 8 | 2 | 1024 | 125 | None |
| 64 | 2 | 4 | 256 | 125 | None |
| 128 | 4 | 4 | 512 | 125 | None |
| 256 | 8 | 4 | 1024 | 125 | None |
| 64 | 2 | 2 | 256 | 125 | None |
| 128 | 4 | 2 | 512 | 125 | None |
| 256 | 8 | 2 | 1024 | 125 | None |
| 64 | 2 | 4 | 256 | 125 | None |
| 128 | 4 | 4 | 512 | 125 | None |
| 256 | 8 | 4 | 1024 | 125 | None |
| 64 | 2 | 2 | 256 | 250 | None |
| 128 | 4 | 2 | 512 | 250 | None |
| 256 | 8 | 2 | 1024 | 250 | None |
| 64 | 2 | 4 | 256 | 250 | None |
| 128 | 4 | 4 | 512 | 250 | None |
| 256 | 8 | 4 | 1024 | 250 | None |
| 64 | 2 | 2 | 256 | 250 | None |
| 128 | 4 | 2 | 512 | 250 | None |
| 256 | 8 | 2 | 1024 | 250 | None |
| 64 | 2 | 4 | 256 | 250 | None |
| 128 | 4 | 4 | 512 | 250 | None |
| 256 | 8 | 4 | 1024 | 250 | None |
| 64 | 2 | 2 | 256 | 125 | [256] |
| 128 | 4 | 2 | 512 | 125 | [256] |
| 256 | 8 | 2 | 1024 | 125 | [256] |
| 64 | 2 | 4 | 256 | 125 | [256] |
| 128 | 4 | 4 | 512 | 125 | [256] |
| 256 | 8 | 4 | 1024 | 125 | [256] |
| 64 | 2 | 2 | 256 | 125 | [256] |
| 128 | 4 | 2 | 512 | 125 | [256] |
| 256 | 8 | 2 | 1024 | 125 | [256] |
| 64 | 2 | 4 | 256 | 125 | [256] |
| 128 | 4 | 4 | 512 | 125 | [256] |
| 256 | 8 | 4 | 1024 | 125 | [256] |
| 64 | 2 | 2 | 256 | 250 | [256] |
| 128 | 4 | 2 | 512 | 250 | [256] |
| 256 | 8 | 2 | 1024 | 250 | [256] |
| 64 | 2 | 4 | 256 | 250 | [256] |
| 128 | 4 | 4 | 512 | 250 | [256] |
| 256 | 8 | 4 | 1024 | 250 | [256] |
| 64 | 2 | 2 | 256 | 250 | [256] |
| 128 | 4 | 2 | 512 | 250 | [256] |
| 256 | 8 | 2 | 1024 | 250 | [256] |
| 64 | 2 | 4 | 256 | 250 | [256] |
| 128 | 4 | 4 | 512 | 250 | [256] |
| 256 | 8 | 4 | 1024 | 250 | [256] |
| **Training hyperparameters** |  |  |  |  |  |
| **Hyperparameter** | **Values** |  |  |  |  |
| Random seeds | 0, 1, 2 |  |  |  |  |
| Batch size | 64 |  |  |  |  |
| Maximum epochs | 50 |  |  |  |  |
| Patience | 20 |  |  |  |  |
| Initial learning rate | 1,00E-03 |  |  |  |  |
| LR scheduler | Cosine annealing + warm-up |  |  |  |  |
| Warm-up epochs | 10 |  |  |  |  |
| Patience | 20 |  |  |  |  |
| Minimum learning rate | 1,00E-08 |  |  |  |  |
| Weight decay | 0, 0.1 |  |  |  |  |
| Dropout, hidden layer | 0, 0.2 |  |  |  |  |
| Data augmentation | Off; low; medium; high |  |  |  |  |
| **Data augmentation** |  |  |  |  |  |
| **Name** | **Noise probability** | **Noise STD** | **Scaling probability** | **Min scale** | **Max scale** |
| Off | 0 |  | 0 |  |  |
| Low | 0,25 | 0,25 | 0,25 | 0,5 | 3 |
| Medium | 0,5 | 0,25 | 0,5 | 0,5 | 3 |
| High | 0,75 | 0,25 | 0,75 | 0,5 | 3 |
| **Hyperparameter search<br>Transformer** |  |  |  |  |  |
| **Models** |  |  |  |  |  |
| **Embedding dimension** | **Attention heads** | **Encoder layers** | **Feed-forward dimension** | **Patch strides** | **FCN hidden layers** |
| 64 | 2 | 2 | 256 | 125 | None |
| 128 | 4 | 2 | 512 | 125 | None |
| 256 | 8 | 2 | 1024 | 125 | None |
| 64 | 2 | 4 | 256 | 125 | None |
| 128 | 4 | 4 | 512 | 125 | None |
| 256 | 8 | 4 | 1024 | 125 | None |
| 64 | 2 | 2 | 256 | 125 | None |
| 128 | 4 | 2 | 512 | 125 | None |
| 256 | 8 | 2 | 1024 | 125 | None |
| 64 | 2 | 4 | 256 | 125 | None |
| 128 | 4 | 4 | 512 | 125 | None |
| 256 | 8 | 4 | 1024 | 125 | None |
| 64 | 2 | 2 | 256 | 250 | None |
| 128 | 4 | 2 | 512 | 250 | None |
| 256 | 8 | 2 | 1024 | 250 | None |
| 64 | 2 | 4 | 256 | 250 | None |
| 128 | 4 | 4 | 512 | 250 | None |
| 256 | 8 | 4 | 1024 | 250 | None |
| 64 | 2 | 2 | 256 | 250 | None |
| 128 | 4 | 2 | 512 | 250 | None |
| 256 | 8 | 2 | 1024 | 250 | None |
| 64 | 2 | 4 | 256 | 250 | None |
| 128 | 4 | 4 | 512 | 250 | None |
| 256 | 8 | 4 | 1024 | 250 | None |
| 64 | 2 | 2 | 256 | 125 | [256] |
| 128 | 4 | 2 | 512 | 125 | [256] |
| 256 | 8 | 2 | 1024 | 125 | [256] |
| 64 | 2 | 4 | 256 | 125 | [256] |
| 128 | 4 | 4 | 512 | 125 | [256] |
| 256 | 8 | 4 | 1024 | 125 | [256] |
| 64 | 2 | 2 | 256 | 125 | [256] |
| 128 | 4 | 2 | 512 | 125 | [256] |
| 256 | 8 | 2 | 1024 | 125 | [256] |
| 64 | 2 | 4 | 256 | 125 | [256] |
| 128 | 4 | 4 | 512 | 125 | [256] |
| 256 | 8 | 4 | 1024 | 125 | [256] |
| 64 | 2 | 2 | 256 | 250 | [256] |
| 128 | 4 | 2 | 512 | 250 | [256] |
| 256 | 8 | 2 | 1024 | 250 | [256] |
| 64 | 2 | 4 | 256 | 250 | [256] |
| 128 | 4 | 4 | 512 | 250 | [256] |
| 256 | 8 | 4 | 1024 | 250 | [256] |
| 64 | 2 | 2 | 256 | 250 | [256] |
| 128 | 4 | 2 | 512 | 250 | [256] |
| 256 | 8 | 2 | 1024 | 250 | [256] |
| 64 | 2 | 4 | 256 | 250 | [256] |
| 128 | 4 | 4 | 512 | 250 | [256] |
| 256 | 8 | 4 | 1024 | 250 | [256] |
| **Training hyperparameters** |  |  |  |  |  |
| **Hyperparameter** | **Values** |  |  |  |  |
| Random seeds | 0, 1, 2 |  |  |  |  |
| Batch size | 64 |  |  |  |  |
| Maximum epochs | 50 |  |  |  |  |
| Patience | 20 |  |  |  |  |
| Initial learning rate | 1,00E-03 |  |  |  |  |
| LR scheduler | Cosine annealing + warm-up |  |  |  |  |
| Warm-up epochs | 10 |  |  |  |  |
| Patience | 20 |  |  |  |  |
| Minimum learning rate | 1,00E-08 |  |  |  |  |
| Weight decay | 0, 0.1 |  |  |  |  |
| Dropout, hidden layer | 0, 0.2 |  |  |  |  |
| Data augmentation | Off; low; medium; high |  |  |  |  |
| **Data augmentation** |  |  |  |  |  |
| **Name** | **Noise probability** | **Noise STD** | **Scaling probability** | **Min scale** | **Max scale** |
| Off | 0 |  | 0 |  |  |
| Low | 0,25 | 0,25 | 0,25 | 0,5 | 3 |
| Medium | 0,5 | 0,25 | 0,5 | 0,5 | 3 |
| High | 0,75 | 0,25 | 0,75 | 0,5 | 3 |
| **Hyperparameter search<br>MLP** |  |  |  |  |  |
| **Models** |  |  |  |  |  |
| **Hidden layers** | **Hidden sizes** |  |  |  |  |
| 0 | - |  |  |  |  |
| 1 | 16 |  |  |  |  |
| 2 | 16 |  |  |  |  |
| 3 | 16 |  |  |  |  |
| 4 | 16 |  |  |  |  |
| 1 | 32 |  |  |  |  |
| 2 | 32 |  |  |  |  |
| 3 | 32 |  |  |  |  |
| 4 | 32 |  |  |  |  |
| 1 | 64 |  |  |  |  |
| 2 | 64 |  |  |  |  |
| 3 | 64 |  |  |  |  |
| 4 | 64 |  |  |  |  |
| **Training hyperparameters** |  |  |  |  |  |
| **Hyperparameter** | **Values** |  |  |  |  |
| Random seeds | 0, 1, 2 |  |  |  |  |
| Batch size | 64 |  |  |  |  |
| Maximum epochs | 50 |  |  |  |  |
| Patience | 20 |  |  |  |  |
| Initial learning rate | 1,00E-02 |  |  |  |  |
| LR scheduler | Cosine annealing + warm-up |  |  |  |  |
| Warm-up epochs | 10 |  |  |  |  |
| Patience | 20 |  |  |  |  |
| Minimum learning rate | 1,00E-08 |  |  |  |  |
| Weight decay | 0, 0.1, 0.01, 0.001 |  |  |  |  |
| Dropout, hidden layer | 0, 0.1, 0.2, 0.3, 0.4 |  |  |  |  |

