## Supplementary Methods for "Extracting generator-specific information from irregular brain field potentials: a comparative study of feature-based and deep learning approaches"

### Metrics definitions

The following expressions describe the mathematical formulation of the measures computed in this work. They are expressed for a given time series $x_n$, $n = 1, \ldots, N$, with sampling frequency $F_s$ and sample mean $\tilde{x} = \frac{1}{N}\sum_{n=1}^{N} x_n$. Unless stated otherwise, *log* denotes the natural logarithm.

#### Temporal measures

*Skewness.*

$$\gamma_1(x) = \frac{\frac{1}{N}\sum_{n=1}^{N}(x_n - \tilde{x})^3}{\left[\frac{1}{N}\sum_{n=1}^{N}(x_n - \tilde{x})^2\right]^{3/2}} \qquad (1)$$

The skewness measures the asymmetry of a distribution. Zero skewness corresponds to a symmetric distribution. For an FP-type time series, it has no simple interpretation as it may be influenced by several parameters concerning individual waveforms or their grouping.

#### Spectral measures

Power spectra are commonly used to highlight the presence of rhythmic oscillations in an FP and their relative power. However, note that any single wave already contains power in a wide power band. Brief periods of oscillatory activity could indeed be present and may strongly determine spectral properties in some segments. However, a given frequency band (e.g., gamma) is not sufficient discriminant feature as it may appear in several generators (Torres et al., 2019; López-Madrona et al., 2020; Hernández-Recio et al., 2023). Here we used global descriptors of the spectral domain (mean, variance, kurtosis and skewness) that may better account for the contribution of non-periodic activity. The power on specific standard bands was also estimated but not included in the group of measures due to low informativeness.

For the spectral mean, variance, kurtosis and skewness, the discrete Fourier transform is calculated:

$$X_k = \sum_{n=1}^{N} x_n \; e^{-i2\pi k(n-1)/N}, \; k = 0, \dots, N-1 \qquad (3)$$

with associated frequencies $f_k = kF_s/N$.

*Power spectral density (PSD)*

For a wide-sense stationary process, the PSD describes the distribution of power across the frequency components of $x_n$ and is the Fourier transform of the autocorrelation function (Wiener–Khinchin theorem):

$$S_{xx}(f) = \int_{-\infty}^{\infty} R_{xx}(\tau)\, e^{-i2\pi f \tau}\, d\tau, \qquad R_{xx}(\tau) = \mathbb{E}[x(t)\, x^{*}(t-\tau)] \qquad (4)$$

For an ergodic process, $R_{xx}$ can be computed as a time average, $R_{xx}(\tau) = \lim_{T\to\infty} \frac{1}{T} \int_{-T/2}^{T/2} x\,(t) x^{*}(t-\tau)\, dt$. From a finite record we use the periodogram estimate

$$\hat{S}_{xx}(f_k) = \frac{1}{NF_s} \, |X_k|^2 \qquad (5)$$

**Measures of entropy**

The entropy of a random variable can be interpreted as the amount of information provided by the observation of a variable. The more random and unpredictable the variable is, the greater its entropy (Rényi, 1961).

For a discrete probability distribution $\{p_i\}_{i=1}^{M}$, Shannon entropy is

$$H_S = -\sum_{i=1}^{M} p_i \log p_i \qquad (6)$$

and Rényi entropy of order $\alpha \neq 1$ is

$H_\alpha = \frac{1}{1-\alpha}\log(\sum_{i=1}^{M} p_i^\alpha)$. (7)

*Wavelet entropies*

A maximal overlap discrete wavelet transform (MODWT) with the Symlet 4 (sym4), with the decomposition level determined automatically from the signal length. Let *$W_j[n]$* be the wavelet coefficients at scale *j*. The relative energy at each level is

$$p_j = \frac{E_j}{E_{\text{tot}}},\ E_j = \sum_n W_j\,[n]^2,\ E_{\text{tot}} = \sum_{j=1}^{J} E_j \qquad (7)$$

The Shannon (*$H_S$*) and Rényi ($H_\alpha$) wavelet entropies are computed from $\{p_j\}_{j=1}^{J}$ with the formulas above, using α=2.

*Spectral entropy*

Let *$P(F_j)$* be the PSD estimated with Welch's method and

$$p_j = \frac{P(f_j)}{\sum_k P(f_k)},\ H_{\text{spec}} = -\sum_j p_j \log p_j \ (\text{optionally divided by } \log N_f) \qquad (8)$$

where *$N_f$* is the number of frequency bins.

*SVD entropy*

Using an embedding of dimension 3 and delay 1, define $X_m = (x_m, x_{m+1}, \dots, x_{m+N-3})^\top$ for *m = 1,2,3,* each of length *N-2*, and the matrix $M = [X_1, X_2, X_3] \in \mathbb{R}^{(N-2)\times 3}$. Let *$M = U\Sigma V^T$* be its singular value decomposition with singular values $\sigma_1$, $\sigma_2$, $\sigma_3$. Then

$$p_i = \frac{\sigma_i}{\sum_{j=1}^{3}\sigma_j},\ H_{\text{SVD}} = -\sum_{i=1}^{3} p_i \log p_i \ (\text{optionally divided by } \log 3) \qquad (9)$$

*Permutation entropy*

Form the vectors $v_n = (x_n, x_{n+1}, x_{n+2})$, $n=1,...N-2$, and map each to its ordinal pattern $\pi \epsilon S_3$. With $p(\pi)$ the relative frequency of pattern $\pi$ among the $N-2$ vectors,

$$H_{\text{per}} = -\frac{1}{\log 3!}\sum_{\pi \in S_3} p\,(\pi)\log p(\pi) \in [0,1] \qquad (10)$$

*Approximate and Sample Entropies*

A detailed discussion of both measures can be found in Delgado-Bonal and Marshak (2019).

**Other complexity measures**

*Negentropy*

For a random variable $X$ with differential entropy $H(X)$, negentropy is

$$J(X) = H(X_G) - H(X) \ \geq 0 \qquad (11)$$

where $X_G$ is a Gaussian variable with the same variance as $X$.

*Higuchi dimension $D_H$*

For an integer scale $k \geq 1$ and an initial point $m = 1,...,k$, consider the subseries $x_m^k = \{x_m, x_{m+k}, x_{m+2k}, ...\}$ and let $N_{m,k} = \lfloor (N-m)/k \rfloor$. Its normalized length is

$$L_m(k) = \frac{N-1}{k^2\, N_{m,k}}\sum_{i=1}^{N_{m,k}} \left| x_{m+ik} - x_{m+(i-1)k} \right| \qquad (14)$$

Then, $D_H$ is computed as:

$$D_H = -\lim_{k\to\infty}\lim_{N\to\infty}\frac{\log\left(\frac{1}{k}\sum_{m=1}^{k} L_m(k)\right)}{\log k} \qquad (15)$$

*Katz dimension $D_K$*

Assuming a unit time step between samples,

$$D_K = \frac{\log(N-1)}{\log(N-1)+\log(d/L)} \qquad (16)$$

where $d$ is the maximum distance from the first point $d = \max_n \sqrt{(n-1)^2 + (x_n - x_1)^2}$ and $L$:

$$L = \sum_{n=2}^{N} \sqrt{1 + (x_n - x_{n-1})^2} \qquad (17)$$
